# Simulation-based inference of epidemiological and phylodynamic models via Neural Posterior Estimation

**DOI:** 10.1101/2025.11.25.690436

**Authors:** Francesco Pinotti, Julien Thézé, Xavier Bailly, Guillaume Fournié

## Abstract

Mathematical models play a central role in understanding and forecasting infectious disease dynamics, but parameter inference is often difficult when likelihoods are intractable. Simulation-based inference (SBI) circumvents this limitation by relying on model simulations. Traditional SBI methods, such as Approximate Bayesian Computation, typically depend on handcrafted summary statistics and scale poorly in high-dimensional settings. Recent advances address these challenges by using flexible statistical and machine-learning surrogates to approximate likelihoods or posterior distributions. Neural Posterior Estimation (NPE) directly learns the posterior from simulated data using neural density estimators, automatically extracting informative representations without ad-hoc summaries. Despite its potential, NPE has been rarely applied in infectious disease epidemiology and phylodynamics. Here, we evaluate NPE for parameter inference in mechanistic models using data from the 2014 Ebola outbreak in Sierra Leone. We consider two case studies: a compartmental transmission model fitted to case and death time series, and a birth–death phylodynamic model fitted to an early Ebola phylogeny. In both settings, NPE yields accurate and reliable posterior estimates. Its amortized nature further enables efficient model calibration and criticism. We conclude that NPE is a flexible and scalable tool for epidemiological and phylodynamic inference, and provide detailed online tutorials illustrating the workflow.

## Introduction

Mathematical models are central tools for infectious disease epidemiology. One of their most appealing features, which was apparent already in the seminal work of Kermack and McKendrick [1], is the ability to explain the shape and tempo of real-world outbreaks from a handful of basic mechanistic processes, e.g. transmission, latency and recovery. Moreover, they enable making epidemic forecasts, evaluate the impact of public health interventions and identify populations at heightened risk of infection [2–4]. Data used to inform transmission models come in various forms, e.g. incident cases, prevalence of infection and immune response, or eventually deaths [5–7]. Phylodynamic models work instead with genetic sequences and phylogenetic trees detailing ancestral relationships among sampled pathogens [8]. Despite the elegance and relative ease of implementation of transmission models, they often pose significant challenges in parameter estimation [9]. These challenges are even more pronounced for the most detailed models, e.g. microsimulations or agent-based models among others, which provide a granular representation of the target population with possibly complex interactions among its constituent elements [10].

The traditional approach to parameter inference relies on the likelihood function *p*(*x*|θ), which expresses the probability of observing data *x* given parameters θ for some underlying model. Inference via maximum likelihood seeks the parameter configurations that maximise *p*(*x*|θ), while Bayesian inference uses the posterior distribution *p*(θ|*x*), which is proportional to the likelihood. As with maximum likelihood, one may optimise the posterior or draw samples θ ∼ *p*(θ|*x*) from it. Both operations can be carried out by repeatedly evaluating the likelihood *p*(*x*|θ) for various θ values and fixed observed data *x*. However, such direct approach is feasible only for the simplest infectious disease models since *p*(*x*|θ) is often unknown or intractable. The likelihood being a function of both the model and the data, its accessibility depends on the relationship between the input θ and the model’s output *x*. Direct evaluation of *p*(θ|*x*) is often difficult or even impossible, particularly for detailed, large-scale epidemiological or phylodynamic models. This limitation has motivated the development of simulation-based inference (SBI) methods, which bypass likelihood computation by learning the relationship between simulated data and parameters.

SBI has now become a popular and reliable alternative to traditional likelihood-based approaches [11]. SBI revolves around the idea that a single simulation that yields pseudo-data *x* given parameters θ, is equivalent to sampling from the likelihood distribution, i.e. *x*∼*p*(*x*|θ). Repeated simulations can thus be used to explore the likelihood function, even when it is intractable or unknown. This property is particularly appealing, because performing simulations is often much easier than explicitly evaluating the likelihood.

Approximate Bayesian computation (ABC) is perhaps the most widely used and one of the earliest methods in the SBI toolkit [12]. The rejection-ABC algorithm couples simulation with a rejection step, which retains only simulated pseudo-data, and the associated parameters, that are sufficiently similar to observed data. Despite their conceptual simplicity and theoretical guarantees, all ABC-like algorithms share some common limitations. First, rejection is not straightforward to calibrate and invariably adds some approximation to posterior estimates [13]. Second, ABC suffers from the curse of dimensionality, which can be partly mitigated by reducing the data to a smaller set of summary statistics *y*. However, most ABC algorithms offer no guidance regarding the choice of the most relevant summary statistics, which is generally informed by expert knowledge. Moreover, replacing the data with summary statistics removes some useful information contained in the data, further exposing inference to a potentially significant and uncontrolled degree of approximation [13].

Recent developments in SBI have focused on alleviating some of these limitations while making inference more sample-efficient and scalable. The general strategy is to replace the likelihood or the posterior distribution with a surrogate function that is easier to approximate and analyse [11]. The synthetic likelihood method, for example, approximates the likelihood over summary statistics with a multivariate normal distribution [14]. Regression-based ABC methods instead assume an explicit parametric relationship between parameters and data that accounts also for heteroskedasticity [15]. Bayesian Optimization for Likelihood-Inference (BOLFI) uses a Gaussian Process model to approximate the relationship among parameters and the discrepancy between model simulations and data [16]. Emulation and history matching is yet another method that uses a statistical model to describe the relationship between simulations and parameters with a focus on efficiently removing ‘implausible’ parameter regions [17].

Neural posterior estimation (NPE) is a recent addition to the SBI toolkit [18,19]. NPE aims to approximate the posterior distribution *p*(θ|*x*) with a flexible function based on one or more neural networks trained on simulated data. NPE thus learns the probabilistic mapping between data and parameters and automatically extracts informative representations of the data that serve as implicit summary statistics, effectively removing the need for handcrafted statistics or ad-hoc distance functions. NPE has been applied to complex inverse problems in physics, evolution, neuroscience and economics [20–28], but its potential in epidemiology and phylodynamics has remained largely unexplored. At present, only a few studies have explored the potential of NPE in the context of infectious disease dynamics [29–31]. Radev et al. [30] introduced *OutbreakFlow*, an NPE estimator that infers parameters of a COVID19 infectious disease model. Chatha et al. [29] applied a Susceptible-Infectious model with heterogeneous rates to carbapenem-resistant Klebsiella pneumoniae prevalence data in a hospital. Wieland et al. [31] considered instead a two-strain stochastic differential equation model. These studies reported a positive performance of NPE with respect to ABC benchmarks and demonstrated the increasing interest towards NPE-based statistical surrogate models. We are not aware of studies that applied NPE to phylodynamic inference of infectious diseases, though the potential benefits of amortized inference algorithms clearly open new avenues in the field [32–34]. While existing studies demonstrate promising performance, adapting NPE to specific problems in infectious disease and phylodynamics modelling is not straightforward.

This manuscript aims to illustrate how NPE can be embedded within a rigorous Bayesian inference workflow in epidemiological and phylodynamic inference. We apply NPE to two real-world case studies that are highly relevant to epidemiological outbreak analyses. Both applications use data from the 2014 Ebola virus (EBOV) outbreak in Sierra Leone. In the first example, we use NPE to fit a Susceptible-Exposed-Infectious-Removed (SEIR) model to a time series of reported cases and deaths. In the second example we fit a phylodynamic model to an early phylogeny of EBOV sequences. We then show how the amortized property of NPE enables effective calibration and evaluation of such models. Finally, we benchmark NPE effectiveness and execution time against Markov Chain Monte Carlo (MCMC), the gold-standard method to sample from posterior distributions, in a simplified theoretical phylodynamic example. We provide a detailed account of all the steps of the NPE workflow, from data processing, to neural architecture selection and statistical checks for detecting potential issues. All examples are accompanied by detailed online tutorials to facilitate implementation in similar and broader epidemiological and phylodynamic modelling applications. Finally, we discuss the implications of these results in the perspective of a robust and efficient use of NPE in real-time epidemiology and phylodynamics.

## Results

### Neural Posterior Estimation

NPE aims to approximate the posterior distribution *p*(θ|*x*) from a collection of simulated samples 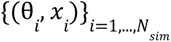 drawn from the joint distribution *p*(*x*, θ). This is achieved by training a *Normalising Flow* (NF) function *q*_φ_(θ|*x*) with hyperparameters φ on these simulations. NFs are highly expressive functions that not only can approximate arbitrary probability distributions with good accuracy, but that are also fast to evaluate and to sample from (see Supplementary Methods for further details on NFs and training) [35]. The trained posterior estimator *q̂*(θ|*x*) should be regarded as a function of both parameters θ and data *x*. This means that if we are interested in performing inference from a specific observation *x_obs_*, we only need to set *x* = *x_obs_* in *q̂*(θ|*x*) to target the corresponding posterior distribution. Please note that NFs are not the only possible choice for *q*_φ_ (θ|*x*) as NPE works with any conditional density estimator.

Once the NPE estimator is trained, switching to a different observation comes with limited computational cost and is nearly immediate, a property called *amortisation*. As we will show later, NPE enables large simulation studies and powerful calibration strategies that would be computationally prohibitive for non-amortised SBI methods.

Another advantage is that NPE can work directly with raw data, bypassing the need to select and calculate summary statistics, as required for standard ABC. Instead, NPE can automatically construct informative summary statistics using a dedicated neural network (see Supplementary Methods).

### 2014 Ebola virus outbreak in Sierra Leone: multivariate time series data

In this section we aimed to infer epidemiological parameters of the 2014 Ebola virus outbreak in Sierra Leone from reported cases and deaths reported in Althaus [36]. We fitted a deterministic transmission model to cases {*c_i_*}_*i*=1_,…,*M* and deaths {*d_i_*}_*i*=1_,…,*M* reported at times {*t_i_*}_*i*=1_,…,*M* (see Methods). This is an example where the likelihood function *p*(*x*|θ) is known analytically, meaning that SBI is not necessary. Nonetheless, it allowed us to compare the estimated posterior *q*(θ|*x*) with the true posterior *p*(θ|*x*), which can be probed via standard MCMC methods.

The transmission model was based on the SEIR model considered in Althaus [36], though our analysis deviates slightly: first, the model was fitted to newly reported rather than cumulative cases. Second, we assumed that reported cases followed a Negative Binomial rather than a Poisson distribution to account for overdispersion in reporting. We inferred 5 parameters: the duration *T*0 between the occurrence of the primary case and the first reported cases, the basic reproduction number *R*_0_, the infection fatality rate *f_deat_*_ℎ_, the impact of public health control interventions *kcontrol* and the shape parameter *s* of the reporting model. The prior distribution is shown in Table S1.

We trained a Masked Autoregressive Flow-based (MAF, a specific type of NF) NPE estimator on *N_sim_* = 50000 draws from *p*(*x*, θ). Each sample *x* was a matrix (2, *M*) where each row represents a different time series of length *M*. The full network architecture is shown in Fig. S1. Once training was done (see learning details in Table S2), the NPE estimator *q̂*(θ|*x*) was extremely fast at drawing posterior samples for any observed data *x_obs_*, and was readily specialized to any observation *x_obs_* (amortization). We first exploited these properties to test the ability of NPE to recover model parameters from 1000 simulated observations. Fig. S2 shows that NPE estimates are accurate almost everywhere in parameter space, except perhaps when the shape parameter *s* is large.

Next, we applied NPE to real data from the 2014 Ebola outbreak in Sierra Leone and compared the NPE estimates with the posterior distribution estimated via MCMC. Fig. 1 shows that the NPE posterior (red) closely matched the true posterior (turquoise), further showcasing the potential of NPE. Our analysis recovered central estimates of *R*_0_, *T*_0_, *f_deat_*_ℎ_ and *k_control_* reported in Althaus [36], while their C.I.s were much narrower. This discrepancy was not specific to the NPE posterior, which we have shown to match the true posterior distribution. Rather, the use of cumulative cases in the original study is likely to have yielded overconfident estimates [37].

**Figure 1:**
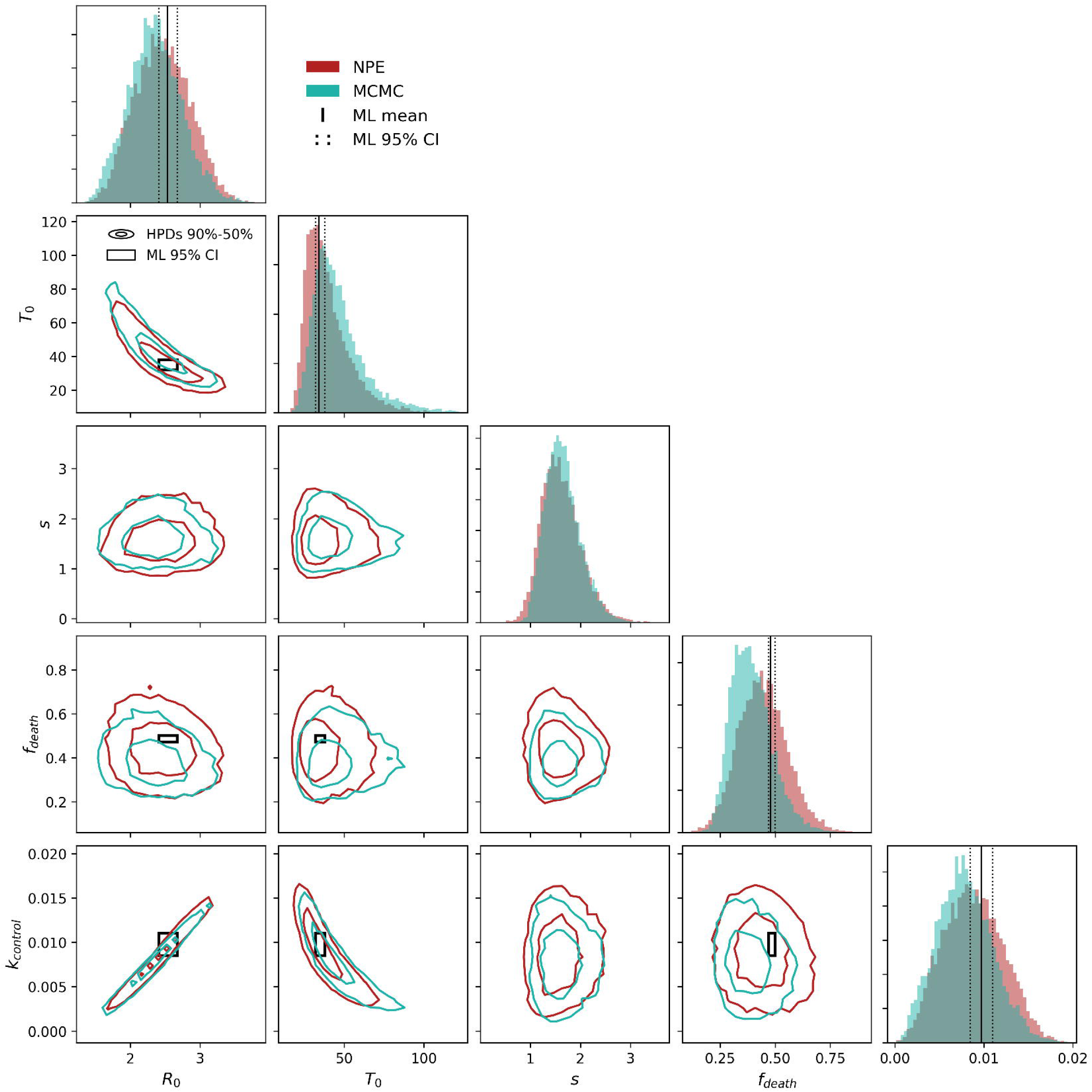
Comparison of NPE and MCMC inference results with time series data. Diagonal plots show posterior marginal distributions obtained using NPE (red) and MCMC (turquoise). Lines represent central estimates (solid) and 95% C.I. (dotted) from Althaus [36]. Please note that there is no reference estimate for *s* as the original model assumed Poisson reporting. Off-diagonal plots show 90% and 50% high posterior density regions (outer and inner circles) obtained from either method. The black rectangle shows the reference estimates. Both distributions are calculated from 10000 posterior samples. MCMC samples were obtained using *Stan* software [38] and the *CmdStanPy* Python interface [39].

Simulations from the posterior predictive distribution were close to the observed time series of reported cases and deaths, demonstrating the quality of our fit (Fig. S3). Amortized inference enabled additional statistically rigorous model checks that would be computationally expensive if not prohibitive with MCMC. Most simulation-based calibration (SBC) methods, for example, require repeated parameter inference on simulated data. Here we considered two popular methods, rank-based SBC and Tests of Accuracy with Random Points (TARP) [40–42], which test whether the estimated posterior displayed the correct coverage properties. Rank-based SBC checks the order statistics (ranks) of true parameters with respect to posterior samples: if the estimated posterior is well calibrated, the ranks are uniformly distributed (Supplementary Methods). Fig. S3 shows that the empirical rank distributions were almost indistinguishable from a uniform distribution for all parameters. By definition, rank-based SBC assesses the coverage property of marginal distributions only. TARP assesses the expected coverage probability of ball-shaped regions in parameter space. Fig. S3 shows that the expected coverage probability estimated via TARP closely matched the credibility level.

The TARP criterion provides a sufficient condition regarding the global correctness of our posterior estimator, on average, over samples drawn from *p*(*x*, θ). While encouraging, however, it is not informative about the local correctness for any specific observation *x_obs_*. Local diagnostics based on classifier 2-sample (C2ST) tests like *l*-C2ST [43] have been recently developed to deal with this question. The rationale behind *l*-C2ST is to train a classifier to distinguish between samples from the true posterior *p*(θ|*x_obs_*) and the estimated posterior *q̂*(θ|*x_obs_*): if the two distributions are indeed similar, the classifier cannot tell them apart. The test proceeded by estimating the deviation of the classifier from chance-level for some observation *xobs* and then constructed a classical p-value. In the case of EBOV case data, we found a small discrepancy with *p* = 0. 21, further supporting the correctness of our estimates.

### 2014 Ebola virus outbreak in Sierra Leone: Phylodynamics inference from sequence data

To further illustrate the relevance of NPE, we analyzed a phylogenetic dataset from the same EBOV outbreak in Sierra Leone. With this purpose, we fitted a mechanistic model to time-scaled phylogenetic trees τ describing the ancestral relationships between 72 published EBOV sequences [44]. Following previous studies [45,46], we used a Birth-Death model with Exposed and Infectious classes (BDEI model). Briefly, this model is a stochastic extension of the SEIR epidemiological model, taking into account the removal of infectious hosts due to the isolation of cases following their active finding and sampling. Contrary to the SEIR model from the previous section, this BDEI model assumed an infinite supply of susceptible hosts, which is only accurate during the early phases of an outbreak. Unknown parameters include *T*_0_, *R*_0_ (as above), the latent period (1/σ), the infectious period (1/µ) and the proportion ρ of sampled infectious hosts. However, as explained in the Methods, these parameters are not fully identifiable without additional information. We thus followed Saulnier et al. [46], which carried a similar analysis on the same data, and set a tight prior on ρ to allow inference of remaining parameters θ = (*T*_0_, *R*_0_, σ, µ).

Birth-Death models are the cornerstone of phylodynamic inference [47]. However, multi-type Birth-Death models such as the BDEI model are not easily tractable, especially when parameters vary in time as in our case (see Methods). Some BEAST packages offer functionalities to fit this kind of model, but they all make numerical approximations to evaluate the likelihood [8,48]. Notable software that can fit a BDEI model to genetic data include Phylodeep [32] and pyBDEI [49], but these do not implement the exact same model–e.g they assume sampling to be always active (*T*_0_ = 0). Therefore, we compared inference results from NPE with a linear regression correction ABC method (ABC-LR, see Supplementary Methods). This is similar to the ABC-LASSO approach used in Saulnier et al. [46] on the same dataset, but uses a Variance Inflation Factor (VIF) criterion to remove highly collinear data features rather than penalizing model complexity, as done in Danesh et al. [50].

In ABC-LR, data *x* consists of a vector of tree summary statistics introduced in Saulnier et al. [46] (see also Table S3). In contrast, NPE can extract informative summaries automatically from a phylogeny τ, provided it is cast into an appropriate vector/matrix format *x*. Here we choose to use the Compact Bijective Ladderized Vectorization (CBLV) method to represent a binary rooted tree τ with *N* = 72 leaves as a (2, *N*) matrix [32]. We stress that the CBLV representation is lossless and considers all the information contained in the tree τ. Both ABC-LR and NPE use a training dataset consisting of 200000 samples from *p*(*x*, θ) (see Table S2 for details on NPE training).

We first applied NPE and ABC-LR to 1000 simulated trees. Fig. S4 shows that both methods perform well at recovering the true parameters. *T*0 and *R*0 were the easiest to recover, while larger values of *d_E_* = 1/σ and *d_I_*= 1/µ are underestimated. Importantly, both NPE and ABC-LR show similar patterns of bias, suggesting that NPE did not introduce any obvious artefact during inference. We observed similar performances in terms of mean relative error and mean relative bias (Fig. S5). NPE slightly outperformed ABC-LR in terms of parameters *R*0, *dE* and *dI*, while ABC-LR performed better on *T*0, especially at low values. Finally, we verified that the NPE posterior estimator is well calibrated by running rank-based SBC and TARP (Fig. S6).

Having validated methods on simulated data, we next applied NPE and ABC-LR to the empirical sequences from the 2014 Ebola virus outbreak in Sierra Leone. In practice, the phylogenetic tree is not directly observed but should instead be inferred from sequence data, ideally jointly with epidemiological mechanistic parameters θ. A common workaround is to first estimate a time-scaled tree via maximum-likelihood (ML) and lsd [51] or TreeTime [52] and then infer demographic parameters in a second step [53]. Here we preferred to use BEAST over ML due to its integrated approach to tree estimation and dating and the tendency of ML approaches to collapse short branches. Crucially, the chosen BEAST model is simpler than the BDEI model (it does not have latency for instance, see Methods) and can be run in a short amount of time on smaller alignments such as this one. Another advantage of BEAST at this stage is its ability to quantify phylogenetic uncertainty by exploring the posterior distribution over trees. For this application we draw 100 posterior trees, which we obtained by further thinning the post-burn-in MCMC samples to ensure topological independence. Larger datasets involving more complicated models (e.g. phylogeographic ones) and rugged posterior tree spaces may require more trees [54]. We then applied NPE and ABC-LR to each posterior tree and aggregated posterior samples in order to account for uncertainty in tree reconstruction. The results are shown in Fig. 2. Both methods yield similar marginal and bivariate posterior distributions. ABC-LR estimates a slightly shorter infectious period and larger *T*_0_ compared to NPE. Inferred marginals align with previous estimates from Stadler et al. [45] although we estimate a shorter latent period *d_E_*. We cannot make a direct comparison with the original MCMC analysis in terms of *T*_0_ since it modelled the start of sampling differently. However, we can make a comparison in terms of the time *T*_1_ to the first reported sample (Fig. S7). NPE and ABC-LR indicate a more recent EBOV introduction than MCMC. This might also explain why the latent period estimated via NPE and ABC-LR is also shorter compared to the MCMC estimate. *l*-C2ST did not flag any issue with calibration when applied to individual trees (*p* > 0. 05 for all 100 posterior trees). The ABC posterior reported in Saulnier et al. [46] differed from our estimates, most likely because the initial analysis relied on a single, maximum-likelihood (ML) tree. We found that NPE and ABC-LR posterior distributions were quite different from each other when considering the maximum clade credibility (MCC) tree (Fig. S8). In particular, NPE estimates were significantly different from those reported in Fig. 2. Upon closer investigation, we found that most clades in the MCC tree were poorly supported and that the MCC tree deviated significantly from both posterior trees and trees drawn from the prior-predictive distribution (Fig. S9). This suggests that such a summary tree is not a good illustration of the tree posterior distribution, which impacts the efficiency of inference of the BDEI model. Also, while ABC-LR more closely recovered the original posterior distribution than NPE (Fig. S9), neither model passed posterior predictive checks (Fig. S10).

**Figure 2:**
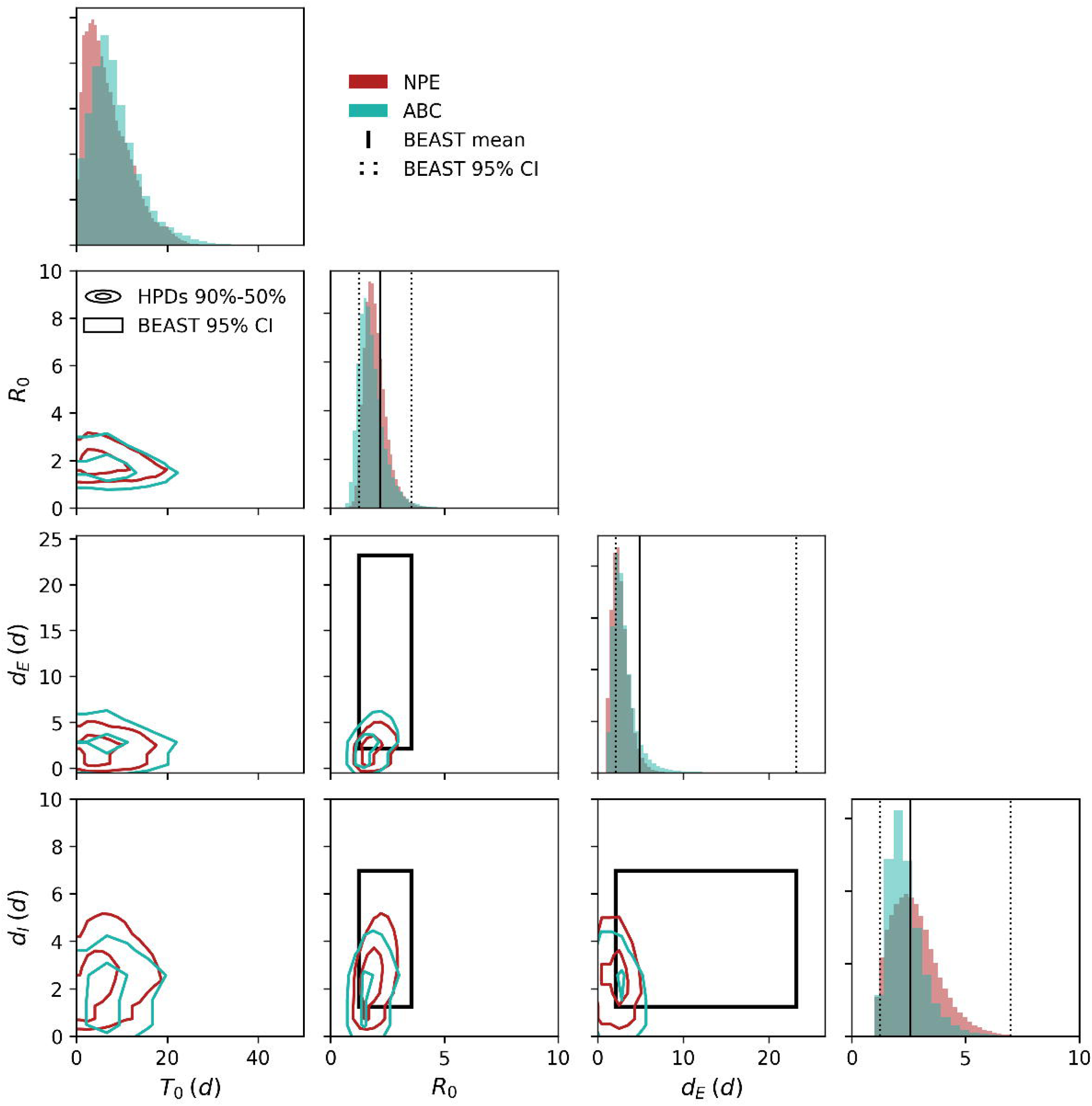
Comparison of NPE and ABC-LR inference results with EBOV sequence data. Diagonal plots show posterior marginal distributions obtained using NPE (red) and ABC-LR (turquoise). Lines represent central estimates (solid) and 95% C.I. (dotted) from Stadler et al. [45]. Off-diagonal plots show 90% and 50% high posterior density regions (outer and inner circles) obtained from either method. The black rectangle shows the reference estimates. We do not show any reference estimate for *T*0 since the original model used a different parameterisation (please check Fig. S7 for further details). Results are based on 10000 posterior samples obtained from 100 posterior trees estimated via BEAST [55].

### Comparing NPE and MCMC performance on a large simulated alignment

To complement the two Ebola case studies, we considered a simplified phylodynamic example designed specifically to assess the performance of NPE and the two-step phylodynamic workflow introduced in the previous section. For this application we simulate an alignment containing 1000 sequences and 2000 characters per sequence using a Birth and Death (BD) tree-generating model and a Jukes-Cantor evolution model [56] with a strict molecular clock. Our aim is to infer *R*_0_ and µ from sequence data (we assume that the sampling ratio is known and equal to ρ = 0. 3). The exact tree, generated under *R*_0_ = 2 and µ = 1 is shown in Fig. 3A. As explained above, our simulation-based pipeline requires (i) a reference table of simulations 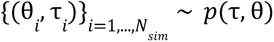 and (ii) a tree estimate τ . Given the relatively large number of sequences, we used ML (IQ-TREE3 [57]) and TreeTime to estimate τ0. This approach yields accurate tree estimates while being much faster than

**Figure 3:**
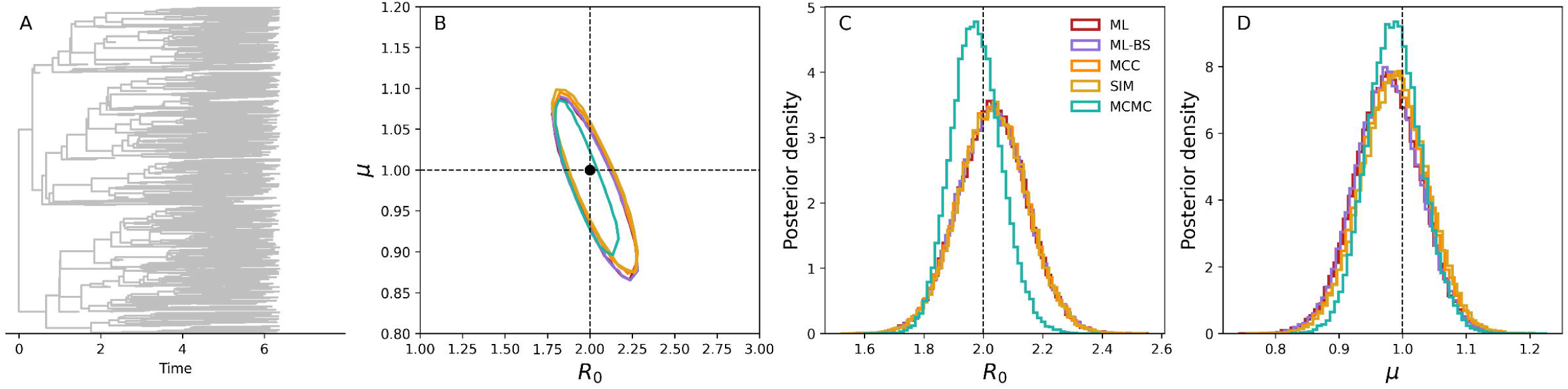
NPE performance on BD tree. (A) Tree simulated under a BD model with *R*_0_= 2, µ = 1 and ρ = 0. 3. (B) 90% highest posterior density regions obtained from 50000 MCMC samples (green) or NPE applied to various trees: the ML tree (red), 50 bootstrap ML trees (purple), the MCC tree from the MCMC analysis (orange) and the original simulated tree (SIM, gold) from panel (A). Marginal distributions for individual parameters are shown in (C,D).

MCMC. We also use IQ-TREE to generate bootstrap trees and quantify phylogenetic uncertainty around the ML solution. We simulated 30000 trees with ReMASTER, which resulted in 22000 valid trees for training. We then used a Mixture Density Network [58] (MDN) as the conditional density estimator for NPE. A single-component MDN enforces a Gaussian posterior whose conditional mean and covariance are jointly estimated via neural networks (see Fig. S1 for the full architecture). While MDNs are less expressive than normalising flows like MAF, they are an appropriate choice in this example where the posterior has a simple elliptical shape (Fig. 3B). Fig. 3B shows various posterior distributions estimated via NPE using different trees. These include the ML tree, a set of 100 bootstrap ML trees, the exact tree from Fig. 3A and the MCC tree calculated from the BEAST analysis. All NPE estimates are similar to each other and recover the true parameters, albeit with a somewhat larger uncertainty than MCMC. Regarding execution time, the entire NPE pipeline took less than 3 hours in total. ML estimation with tip dating, tree simulation, tree encoding and NPE training took 0.06, 2.03, 0.26 and 0.25 hours, respectively. Tree simulation and encoding were parallelised over 6 cores. In contrast, 10^8^ MCMC iterations took 10.81 hours when using the Beagle library [59]. BEAST yielded effective sample sizes around 50000-70000 after accounting for a 10% burning and a 1-in-1000 thinning. Please note that MCMC must be run for longer to accumulate independent posterior samples, while NPE can draw as many independent samples as desired with no cost in terms of computation time.

## Discussion

We showed that Neural Posterior Estimation yields reliable Bayesian estimates in the context of real outbreak analyses characterised by noisy, hierarchical and multivariate data. We considered two models related to the 2014 EBOV outbreak in Sierra Leone where data consisted either of reported cases or genetic sequences. In the former case, we showed that NPE nearly recovered the true posterior distribution. In the second example the likelihood function was not available and we tested NPE against regression ABC, another popular simulation-based inference method. Both methods yielded similar estimates on both simulated and posterior trees, which hints at NPE as a valid and competitive simulation-based inference method.

From a computational perspective, simulation and training steps are the main bottlenecks. We showed that NPE and ABC-LR are both efficient in terms of simulation requirements, using only 50000, 200000 and 22000 simulations in the SEIR, BDEI and BD applications, respectively. Other SBI algorithms, e.g. rejection ABC or sequential variants of ABC, are typically much less efficient and may require millions of simulations [60]. Training time depends mostly on neural network size, the number of training simulations and the training settings. Here we were able to complete training on an NVIDIA A40 gpu in about 1 hour in both SEIR and BDEI applications, and in less than 15 minutes in the BD case. After training, the NPE estimator is extremely fast to sample from (it took around 3s to draw 1000000 samples on our server) and can be readily applied to new observations thanks to its amortised nature.

Both regression ABC and NPE aim to approximate the conditional density *p*(θ|*x*). Conceptually, they each try to model how θ varies with data *x* or with derived summary statistics *y*(*x*): our version of regression ABC does this locally via a linear heteroskedastic correction near the observed summaries [15], while NPE does it globally with a flexible neural density model. Phylodeep [32] also uses neural networks to recover mechanistic parameters from raw trees via regression. However, the method focuses on retrieving point estimates rather than a joint posterior distribution, although confidence intervals can still be obtained.

NPE can use a dedicated neural network to automatically learn summary statistics. This is a considerable advantage over ABC, since selecting appropriate summary statistics requires substantial care. Nonetheless, one must still choose an appropriate neural network for feature extraction, which is not a straightforward task either. One strategy would be to train multiple candidate architectures and choose the one with the smallest error on validation data. Smaller architectures require less data but risk underfitting, while more complex and larger architectures risk overfitting. Overfitting means that the inferred posterior estimator yields good results on training data but fails to generalise to new data points, e.g. those used for validation. Here we did not perform a systematic investigation as it was beyond the scope of our study, and relied on heuristic considerations and validation error as guiding principles. A general recommendation is to adopt networks with the appropriate inductive biases, i.e. networks that are wired to recognise and recover the correlational structure of the data. For example, convolutional neural networks can learn short-scale correlations that characterise, for example, image data [61]. Both our examples make use of stacks of convolutional and fully-connected layers (Fig. S1), which can tackle data that are inherently hierarchical, as expected from time series and phylogenetic trees. The feature extractor in the phylodynamics examples is indeed based on published networks such as Phylodeep [32]. Other types of networks may also be envisioned, such as recurrent neural networks for temporal data, graph neural networks and transformers [62–64]. Transformers have been used for example to process multi-sequence alignments [33,65] and can allow parameter inference without the need to estimate the tree at all [33]. It should be stressed that similar choices must be made about the conditional density estimator (here we used MAF and Mixed Density Networks), the number and size of layers as well as estimator-specific hyperparameters. Ultimately, users should select architectures whose complexity is commensurate with the number of training simulations.

NPE directly targets the posterior distribution *p*(θ|*x*), which has the advantage of making sampling as fast as a single forward pass of the corresponding neural network. Other neural-based methods like Neural Likelihood Estimation (NLE) and Neural Ratio Estimation (NRE) target instead the likelihood *p*(*x*|θ) and the likelihood-to-evidence ratio *p*(*x*|θ)/*p*(*x*), respectively [66,67]. These targets might be easier to learn than the posterior, but sampling still requires a round of MCMC. Moreover, NLE cannot use a feature extractor, limiting its scalability to large data. Sequential versions of NPE, NLE and NRE can increase the accuracy of these algorithms on specific observations with less simulations, but the resulting posterior estimators are not amortized [19,66–68]. Finally, NPE methods that are based on diffusion models and continuous NFs have also been proposed [69,70]. These methods put less constraints on the network architecture than discrete NFs (like the MAF considered here), which can only accommodate invertible functions with a simple Jacobian structure as building blocks.

We showed that NPE can be easily incorporated within a principled Bayesian framework [71]. Amortisation paves the way for a range of simulation-based calibration tools that may be too time-consuming with other inference methods [72]. Rank-based SBC and related methods are relatively simple checks that can detect patterns of bias, overconfidence or underconfidence [40,41]. TARP works similarly, but provides necessary and sufficient conditions for a globally accurate posterior estimation [42]. Local methods like *l*-C2ST can help detect issues with specific problem instances [43].

As any other inference method, NPE has some limitations. One issue is that NPE may not perform robustly in some cases, e.g. when the model is unable to generate simulations that resemble the data. In this case, NPE extrapolates beyond the training data. We have seen this phenomenon in the context of the MCC EBOV tree, which was quite different from training trees. On one hand, this example warns against using a single summary tree when phylogenetic uncertainty is high [73]. On the other hand, it underlies the importance of verifying model adequacy via, e.g., prior predictive checks. Analogously, NPE may also struggle to estimate the posterior distribution when some observations lie far from training points [74]. Increasing the size of the training set can alleviate this issue, at the cost of increased simulation and training times.

This study considers relatively simple but relevant epidemic models that are commonly found in the literature. We believe that NPE will be even more useful in the context of more complex models, e.g. agent-based models that are often regarded as stochastic black-boxes. Indeed, NPE was shown to enable inference in the case of a model of social dynamics [27]. Finally, we note that all our analyses focus on a past outbreak and are hence retrospective in nature. Nonetheless, we believe that NPE may also facilitate model calibration in real time. This means dealing with rapidly unfolding outbreaks where data (cases, sequences) accumulate on a daily basis and model predictions must be updated with the same frequency. One potential strategy would be to train an NF once on simulations of varying sizes and exploit the amortized property to rapidly generate posterior estimates without any need for further training.

## Methods

### EBOV Time series data

Cumulative EBOV cases and deaths reported during the 2014 outbreak in Sierra Leone were obtained from Althaus [36]. *C_i_* (*D_i_*) denotes cumulative cases (deaths) reported up to time *t_i_*, where *i* = 1, …, *M*’, *M*’ is the length of the original time series and times *t*_1_, *t_M_*’ correspond to dates 27/05/2014 and 20/08/2014, respectively. We then calculated newly reported cases and deaths as *c*_1_= *C*_1_, *d*_1_ = *D*_1_ and *c_i_*= *C_i_* − *C_i_*_−1_, *d_i_*= *D_i_* − *D_i_*_−1_ for *i* > 1, skipping intermediate time points for which any of these differences were negative. In this case, we search for the first time point *j* > *i* for which both *C_j_* ≥ *C_i_* and *D_j_* ≥ *D_i_*. The resulting time series *c_i_*, *d_i_* are therefore shorter than the original data (*i* = 1, …, *M* < *M*’). Cases and deaths are log-transformed according to log(1 + *x*) before being passed to neural networks. Moreover, the neural network z-scores each input separately.

### EBOV sequence data

We used 72 full-length EBOV sequences collected from 26/05/2014 to 18/06/2014 in Sierra Leone and originally reported in Gire et al. [44].

### Time series SEIR epidemic model

The numbers of susceptibles *S*, exposed *E*, infectious *I* and recovered *R* evolve according to the following dynamical equations:

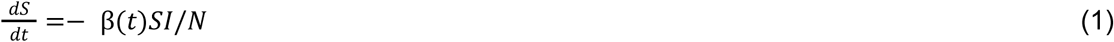

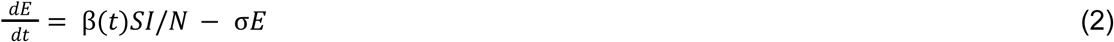

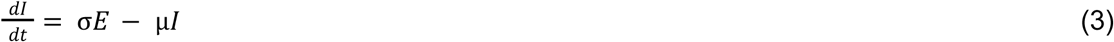

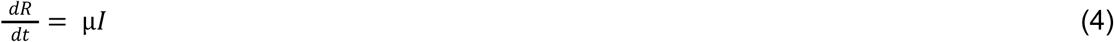

Where *N* = 10_6_ denotes the total population size, σ^−1^ = 5. 3 days is the latent period and µ^−1^ = 5. 61 days is the infectious period. The transmission rate is defined as 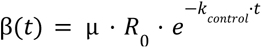, where parameters *R*_0_, *k_control_* are to be estimated. The parameter *k_control_* > 0 quantifies the effectiveness of control measures: the larger *k_control_*, the faster the decay of the transmission rate. Infected individuals have a probability *f_death_* to die. The outbreak begins with a single infectious individual at time *t* = 0, but cases are only reported after a delay *T*_0_ to be estimated.

We assume a negative binomial observation model for both cases and deaths. More in detail, the expected numbers of cases *c_i_* and deaths *d_i_* reported between *t_i_*_−1_ and *t_i_* follow a negative binomial distribution with means 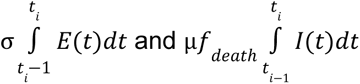, respectively. The shape parameter *s* is the same for both data sources and has to be estimated. This observation model assumes that all cases are reported on average, but allows some uncertainty around the expected number of cases. The full likelihood function is then given by:

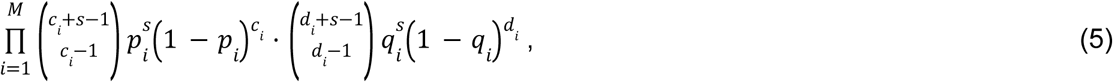

where 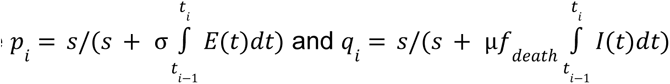

We retain only simulations where total cases fall between 100 and 1000000.

### Phylogenetic analysis of EBOV sequences

We used BEAST v2.7.8 [55] to infer and date a phylogeny of available genetic sequences. We used a Birth-Death model as the tree prior, a HKY substitution model with gamma-distributed heterogeneity and 4 rate categories [75,76]. We implemented the Birth-Death model using the BDMM-Prime BEAST package [48,77], setting the sampling ratio to 0 before the oldest sample. We assume a strict molecular clock with a fixed rate 0.001984 _−1_ [45]. We use the same prior specifications as in the BD 1 model in Stadler et al. [45] for all parameters but the *origin* parameter (the time between the earliest sample and the start of the outbreak), whose prior we assume to be uniform between 0 and 1000 days for computational convenience. We run a single MCMC chain with 20000000 steps. We assessed convergence using Tracer v1.7.2 [78] and checked that the effective sample size of each estimated parameter was above 200. We obtained the MCC tree with TreeAnnotator v2.7.7 [79].

### Birth-death model with exposed individuals (BDEI model)

According to this model, an infectious individual generates another infection with rate β = *R*_0_· µ or is removed with rate µ due to recovery, death or sampling. The sampling ratio ρ denotes the probability that a removed infection is also sampled. Newly infected individuals enter the exposed compartment, they cannot be sampled and become infectious with rate σ. Moreover, we assume that sampling starts only after *t* > *T*_0_ and set the sampling ratio to 0 for *t* < *T*_0_. A simulation stops when exactly 72 samples have been collected, yielding a time-scaled phylogeny determined by the full transmission chain. We reject simulations that fizzle out before reaching the desired number of samples or produce more than 50000 cases. We do not try to infer the sampling ratio ρ since it is well known that this parameter cannot be identified jointly with µ and *R*0 [80]. Following Saulnier et al. [46], we assume that ρ is approximately around 0.7 but still allow for some variability around this value; in practice, we draw ρ from a narrow prior distribution centered around 0.7 (see Table S4) and use these values to generate training data. We run BDEI simulations using the ReMASTER [81] package in BEAST v2.7.8.

### Phylodynamic simulation and analysis of the BD model

In order to illustrate further computational cost and the properties of NPE approaches, we designed a simplified benchmark to complement the Ebola case studies. We assumed a simplified version of the BDEI model with no latency and no delayed sampling (*T*_0_ = 0). We use ReMASTER to simulate an “observed” alignment of 1000 sequences and 2000 characters under *R*_0_ = 2, µ = 1 and ρ = 0. 3, as well as pairs 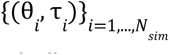 to train NPE. where τ denotes a time-scaled tree. Each site in the simulated alignment evolves independently according to a Jukes-Cantor model [56] and we impose a strict molecular clock with rate 0.05. This ensures that the alignment is informative about the phylogeny. We estimate an ML tree using IQ-TREE3 [57] with options -m JC -bb 1000 -wbt to ensure a Jukes-Cantor evolution process and the simultaneous generation of 1000 bootstrap trees via the ultrafast bootstrap method [82]. We date the ML and bootstrap trees with the TreeTime v0.11.4 Python module [52] under a strict molecular clock. The BEAST MCMC analysis uses the very same generative model for inference. We assign uniform priors to parameters *R*0 and µ, which we assume to range between 1 and 5 and between 0 and 5, respectively. The sampling ratio ρ is always set to 0.3 for the sake of identifiability.

### Ethics statement

The study did not involve the collection of new data from human participants or animals. In accordance with institutional and national guidelines, ethical approval and informed consent were not required.

### Data accessibility

All data used in this study were obtained from publicly available sources. Time series of EBOV case counts and deaths were obtained from Althaus [36]. EBOV sequences were obtained from Gire et al. [44]. Code, data and relevant tutorials are available on GitHub at https://forge.inrae.fr/francesco.pinotti/SBI-EPI-PHYLO-tutorial. The final version of the GitHub repository has been archived on Zenodo (https://zenodo.org/records/20306569) [83]. Additional methodological details and figures are available as electronic supplementary material.

### Authors’ contributions

F.P.: conceptualization, formal analysis, investigation, methodology, project administration, software, visualization, funding acquisition, writing—original draft, writing—review and editing; J.T.: conceptualization, methodology, supervision, writing—review and editing; X.B.: conceptualization, methodology, supervision, writing—review and editing; G.F.: conceptualization, methodology, funding acquisition, supervision, writing—review and editing. All authors gave final approval for publication and agreed to be held accountable for the work performed therein.

### Declaration of AI use

We have not used AI-assisted technologies in creating this article.

## Acknowledgements

We thank David Abrial for reviewing the code and helping us set up an online code repository.

