## Supplementary Methods for "Simulation-based inference of epidemiological and phylodynamic models via Neural Posterior Estimation"

### Electronic supplementary material for the manuscript "Simulation-based inference of epidemiological and phylodynamic models via Neural Posterior Estimation"

Francesco Pinotti[1,2], Julien Thézé[1,2], Xavier Bailly[1,2], Guillaume Fournié[1,2,3]

[1]Université de Lyon, INRAE, VetAgro Sup, UMR EPIA, Clermont-Auvergne-Rhône-Alpes, Lyon, France
[2]Université Clermont Auvergne, INRAE, VetAgro Sup, UMR EPIA, Saint-Genès-Champanelle, France
[3]Department of Pathobiology and Population Sciences, Royal Veterinary College, London, United Kingdom

Published in Proceedings of the Royal Society B: Biological Sciences. doi.org/10.1098/rspb.2026.1059

#### Supplementary methods

**Further details on Neural Posterior Estimation and Normalising Flows.** Mathematically, the NPE estimator $\widehat{q}(\theta|x) = q_{\widehat{\varphi}}(\theta|x)$ corresponds to the solution $\widehat{\varphi}$ of the following optimisation problem:

$$\widehat{\varphi} = argmin_{\varphi} - \sum_{i=1}^{N_{sim}} \log q_{\varphi}(\theta_i|x_i) . \qquad (S1)$$

As explained in the Results section, we choose $q_{\varphi}(\theta|x)$ to be a *Normalising Flow*. An NF is defined as an invertible differentiable transformation $V = F_{\varphi}(U)$ with parameters $\varphi$ mapping a 'simple' random variable $U$ with density $p_U(u)$, e.g. a multivariate normal or uniform, into a new variable $V$ with density $p_V(v)$. These densities are related by the change of variable formula:

$$p_V(v) = p_U(F_{\varphi}^{-1}(v)) \left|\det \partial_u F_{\varphi}^{-1}(v)\right| , \qquad (S2)$$

where $\partial_u F_{\varphi}^{-1}(v)$ is the Jacobian of the inverse transformation $F_{\varphi}^{-1}$ that maps $V$ back into $U$. Under appropriate conditions, NFs enable both fast sampling and evaluation of the target distribution $q_V$. Sampling amounts to calculating $v = F_{\varphi}(u)$ with $u \sim p_U$, while density evaluation exploits the change of variable formula (Eq. S2). Highly expressive NFs that are also computationally convenient can be constructed by stacking multiple simpler transformations with a triangular Jacobian.

Here we consider *Masked Autoregressive Flows* (MAF) as the NF of choice for NPE inference [1]. MAF stacks multiple Masked Autoencoder for Density Estimation (MADE) layers. The MADE is a feed-forward neural network with a masked weight matrix. Masking refers to specific weights being set to 0 in order to enforce autoregressive dependencies among network outputs.

We also consider Mixture Density Networks (MDNs) [2] as yet another conditional density estimator. A Gaussian MDN is a mixture of $K$ Gaussian distributions whose conditional means and covariance functions are estimated via neural networks. Increasing $K$ makes the MDN more expressive, while setting $K = 1$ enforces a multivariate Gaussian posterior distribution.

The target distribution in the NPE context is $q_V = q_\varphi(\theta|x)$, i.e. the approximate posterior. Technically speaking, the target variable $V$ is just the parameters $\theta$, while the data $x$ represents some side-information on which the conditional density estimator is conditioned to. Setting $x = x_{obs}$ in $q_\varphi(\theta|x)$ for some observed data $x_{obs}$ yields an estimate of the posterior $p(\theta|x_{obs})$. This means that inference via NPE is *amortized*, i.e. it can be readily performed on any dataset $x_{obs}$by simply plugging $x_{obs}$ in $q_\varphi(\theta|x)$.

NPE automates the construction of summary statistics from data $x$ by training a second neural network parameterised by $\phi$ that calculates an embedding $y = G_\phi(x)$ that is then fed to $F_\varphi$. Crucially, NPE learns $G_\phi$ and $F_\varphi$ in a joint fashion by solving the following problem:

$$(\widehat{\varphi}, \widehat{\phi}) = argmin_{\varphi,\phi} \; - \sum_{i=1}^{N_{sim}} \log \, q_\varphi(\theta_i|G_\phi(x_i)) \, , \qquad \text{(S3)}$$

Minimising the loss function expressed in Eq. S1 and Eq. S3 is a non-convex optimization problem with no analytical solution. Nonetheless, good solutions can be found by means of heuristic algorithms such as stochastic gradient descent.

**NPE implementation and training.** We used the NPE algorithm implemented in the *sbi* Python module [3]**.** *sbi* provides a friendly interface and implements many neural-based conditional density estimation algorithms and SBC metrics discussed in this manuscript. It relies on the popular *pytorch* module [4] to build neural networks and can run on GPUs to speedup training.

**Simulation-based calibration.** SBC methods aim to assess the accuracy of posterior estimates (including MCMC) without directly accessing the likelihood function $p(x|\theta)$, whether the latter is available or not. All SBC methods discussed here are based on $N_C$ calibration samples $\{(\theta_i, x_i)\}_{i=1,...,N_C}$ drawn from the joint distribution $p(x, \theta) = p(x|\theta)p(\theta)$, which is straightforward to simulate from under our assumptions.

SBC methods may assess local or global accuracy of the estimated posterior depending on whether they refer to a specific observation or not. Rank-based SBC and TARP belong to the latter group. Rank-based SBC generates $N_S$ posterior samples $\theta_i^j \sim \hat{q}(\theta|x_i)$ , $j = 1, ..., N_S$, where $x_i$ is the $i$-th calibration observation ($i = 1, ..., N_C$) simulated given $\theta = \theta_i$. It then proceeds to measure the rank $r_i$ of $\theta_i$ among values $\{\theta_i^j\}_{j=1,...,N_S}$. If $\theta$ is one-dimensional, $r_i = \sum_{j=1}^{N_S} I(\theta_i < \theta_i^j)$ and ranges between 0 and $N_S$. The rationale behind rank-based SBC is that if $\hat{q}(\theta|x_i)$ truly matches the posterior $p(\theta|x_i)$, then $\theta_i$ and $\theta_i^j$ are i.i.d. variables and the ranks $r_i$ should be uniformly distributed between 0 and $N_S$. Any deviation from the uniform distribution suggests that the posterior estimator $\hat{q}(\theta|x)$ is not perfectly calibrated and may thus expose signs of bias, overconfidence and/or underconfidence. If $\theta$ is multivariate, rank-based SBC can be applied to individual components separately.

TARP is another global calibration method that looks at the expected coverage of the posterior estimator. The TARP criterion provides a sufficient condition to assess the global accuracy of $\hat{q}(\theta|x)$, whereas other tests such as rank-based SBC only provide necessary but not sufficient conditions. As in rank-based SBC, TARP generates samples $\theta_i^j$ , $j = 1, ..., N_S$ from $\hat{q}(\theta|x_i)$ for every sample $(\theta_i, x_i)$. It then generates a reference point $\theta_i^r$ from an arbitrary reference point sampling distribution (here we use the prior distribution to make these proposals) and calculates the fraction $f_i$ of samples $\theta_i^j$ within a hyper-ball centered around $\theta_i^r$ and extending up to $\theta_i$. Theory indicates that for well-calibrated posteriors, the distribution of fractions $f_i$ is uniform, indicating that the expected coverage probability matches the credibility level. The corresponding cumulative density should then look like a straight line with unit slope. As in rank-based SBC, any deviations from the uniform distribution may be indicative of patterns of bias, overconfidence, underconfidence, etc.

*l*-C2ST is a local calibration method. As explained in the Results section, *l*-C2ST trains a classifier to distinguish samples $(\theta, x) \sim \hat{q}(\theta|x) \cdot p(x)$ (labelled $l = 0$) from samples $(\theta, x) \sim p(x, \theta)$ (labelled $l = 1$). In practice, we recycle $N_C$ existing draws $(\theta_i, x_i, l_i = 0)$ from $p(x, \theta)$ to generate data points $(\hat{\theta}_i, x_i, l_i = 1)$ where $\hat{\theta}_i \sim \hat{q}(\theta|x_i)$. Note that this operation does not require much overhead if inference is amortized. We then train a binary classifier on this augmented sample with $2N_C$ data points and let $h(x, \theta)$ denote the classifier probability of class $l = 1$ for observation $(\theta, x)$. Let us now consider a specific observation $x_{obs}$ and samples $\{\theta_k\}_{k=1,...,N_S}$ from the estimated posterior $\hat{q}(\theta|x_{obs})$. If $\hat{q}(\theta|x_{obs}) = p(\theta|x_{obs})$, an ideal classifier (in the sense of Bayes-optimal) would not be able to tell whether these samples come from one distribution or the other and $h(x_{obs}, \theta_k) = 0.5$. On the contrary, any deviation from chance-level predictions indicates some difference between estimated and true posterior. To measure this discrepancy, *l*-C2ST calculates the test statistic

$\hat{t}(x_{obs}) = N_S^{-1} \sum_{k=1}^{N_S} (h(x_{obs}, \theta_k) - 0.5)^2$. The measured discrepancy $\hat{t}(x_{obs})$ may be compatible with noise and not reflect a genuine difference between $\hat{q}(\theta|x_{obs})$ and $p(\theta|x_{obs})$. To tease these situations apart, $\hat{t}(x_{obs})$ must be compared against a null distribution of $\hat{t}$ values obtained by training additional classifiers on data with permuted labels. This procedures yields a p-value that quantifies the probability of observing a discrepancy equal to $\hat{t}(x_{obs})$ or larger under the null hypothesis that $\hat{q}(\theta|x_{obs}) = p(\theta|x_{obs})$. In this work we use $N_c = 5000$ and $N_c = 25000$ calibration samples in the SEIR and BDEI examples, respectively.

**ABC-regression with VIF selection of summary statistics.** Following Saulnier et al. [5], we use an ABC regression approach to fit the BDEI model to a given phylogeny. First, we generate $N_{sim} = 200000$ samples $\{(\theta_i, \tau_i)\}_{i=1,\dots,N_{sim}}$ from $p(\tau, \theta)$ and calculate a vector of summary statistics $x(\tau)$ from each tree $\tau$. We use the same summary statistics proposed in [5], with a few minor modifications (a full list of summary statistics is reported in Table S3). We reject simulations where any summary statistics is undefined and divide each summary statistics by its mean absolute deviance so that all summaries share the same scale. Second, we remove all summary statistics with zero variance and those that are highly collinear. Specifically, we remove all summary statistics with a VIF above the recommended threshold of 10. We adopt a greedy strategy, removing the summary statistics with the largest VIF, recalculating VIFs for the remaining statistics and repeating this procedure until no VIF is larger than 10. This procedure results in a reference table $X$ containing $N_{sim}$ rows and $N_{ss} = 31$ columns denoting the remaining summary statistics. Third, given an observed tree $\tau_{obs}$ with summaries $x_{obs}$, we select the $N_{abc} = 10000$ simulations with the smallest euclidean distance $d_i = ||x_i - x_{obs}||$ from $x_{obs}$ and use these in the subsequent regression step. Previous work showed that ABC-regression is robust with respect to the choice of $N_{abc}$, although enough samples should be included to estimate regression parameters reliably [5]. ABC-regression assumes a parametric relationship $\theta_i = m(x_i) + \sigma(x_i)\varsigma$ between parameters and summary statistics, where $m(x)$ is the regression function, $\sigma(x)$ is the square root of the conditional variance of $\theta$ given $x$ accounting for stochasticity, and $\varsigma$ is the residual. Once estimates $\widehat{m}(x)$ and $\widehat{\sigma}(x)$ are available, samples $\theta_i^c$ from the approximate posterior are obtained by correcting samples $\theta_i$ according to $\theta_i^c = \widehat{m}(x_{obs}) + \frac{\widehat{\sigma}(x_{obs})}{\widehat{\sigma}(x_i)} (\theta_i - \widehat{m}(x_i))$. Here we assume that both $m(x)$ and $\sigma(x)$ are linear functions. In practice, these functions are estimated in two steps. First, $m(x)$ is estimated via weighted least squares, i.e. by minimizing the weighted sum of residuals:

$$\sum_{i=1}^{N_{abc}} w_i \cdot (\theta_i - m(x_i))^2, \tag{S4}$$

where weights $w_i$ penalise simulations that are far from $x_{obs}$. These weights are calculated $w_i = 1 - d_i^2/(d_{max} + 0.01)^2$, where $d_{max}$ is the largest euclidean distance among the simulations in the reference table. Next, $\log \sigma^2(x)$ is fitted to the logarithm of the empirical residuals $\widehat{\varepsilon}_i = \theta_i - \widehat{m}(x_i)$ by minimising the quantity:

$$\sum_{i=1}^{N_{abc}} w_i \cdot (\log \widehat{\varepsilon}_i^2 - \log \sigma^2(x_i))^2 . \quad \text{(S5)}$$

We apply the regression-based correction after mapping model parameters to the unbounded domain via a transformation $\theta' = f(\theta)$. The transformation is either a logit or a logarithm function depending on parameter support and prior range bounds. In the phylodynamics example we apply a logarithm transformation to $R_0$ and a logit transformation to the remaining parameters. This transformation guarantees that the corrected parameters do not lie outside the desired range.

#### Supplementary tables

**Table S1**: **Prior distributions for the SEIR model**.

| **Parameter** | **Description** | **Prior** |
|---|---|---|
| $R_0$ | Basic reproduction number. | Uniform(1,4) |
| $T_0$ | Delay between the occurrence of the primary case and the first cases being reported. | Uniform(0 d,120 d) |
| $s$ | Shape of Negative Binomial reporting distribution. | Uniform(0.1,20) |
| $f_{death}$ | Infection fatality rate. | Uniform(0.1,0.9) |
| $k_{control}$ | Impact of control measures | Uniform(0,0.02) |

**Table S2**: **Conditional density estimators' architectural details**

| **Parameter** | **SEIR model** | **BDEI model** | **BD model** |
|---|---|---|---|
| Conditional density estimator | MAF | MAF | MDN |
| Normalise input features? | Yes | No | No |
| Number of layers | 5 | 5 | 25 |

| | | | |
|---|---|---|---|
| Number of hidden units in each layer | 50 | 256 | 32 |
| Dropout | 0.05 | 0 | 0 |
| Optimizer | Adam | Adam | Adam |
| Learning rate | 0.0005 | 0.0005 | 0.0005 |
| Proportion of training samples dedicated to estimate validation error | 0.1 | 0.02 | 0.02 |
| Number of iterations with no improvement in terms of validation loss before stopping | 50 | 30 | 50 |

**Table S3**: summary statistics used in ABC.

| **Name** | **Shortened name** | **Description** |
|---|---|---|
| Maximum temporal leaf-root distance | max_H | Largest temporal distance between the root and the leaves. |
| Minimum temporal leaf-root distance | min_H | Smallest temporal distance between the root and the leaves. |
| Mean branch length | a_BL_mean | Mean branch length calculated from all branches. |
| Median branch length | a_BL_median | Median branch length calculated from all branches. |
| Branch length standard deviation | a_BL_std | Standard deviation of branch lengths calculated from all branch lengths |
| Mean branch length (leaves) | e_BL_mean | Mean branch length calculated from external branches only. |
| Median branch length (leaves) | e_BL_median | Median branch length calculated from external branches only. |
| Branch length standard deviation (leaves) | e_BL_std | Standard deviation of branch lengths calculated from external branches only. |
| Mean branch length (internal) | i_BL_mean.x | Mean branch length calculated from internal branches only. This is calculated over 3 consecutive time periods*. |

| | | |
|---|---|---|
| Median branch length (internal) | i_BL_median.x | Median branch length calculated from internal branches only. This is calculated over 3 consecutive time periods*. |
| Branch length standard deviation (internal) | i_BL_std.x | Standard deviation of branch lengths calculated from internal branches only. This is calculated over 3 consecutive time periods*. |
| Mean branch length ratio | ie_BL_mean.x | Ratio of mean length of internal and external branches. This is calculated over 3 consecutive time periods*. |
| Median branch length ratio | ie_BL_median.x | Ratio of median length of internal and external branches. This is calculated over 3 consecutive time periods*. |
| Branch length standard deviation ratio | ie_BL_std.x | Ratio of the standard deviation of internal and external branch lengths. This is calculated over 3 consecutive time periods*. |
| Maximum number of extant lineages | max_L | Maximum number of extant lineages at any time as seen in the LTT** plot. |
| Timing of maximum number of extant lineages | t_max_L | Time corresponding to the maximum number of extant lineages as seen in the LTT plot. |
| LTT Slope 1 | slope_1 | Slope of the line connecting the origin of the LTT plot and the point in the LTT plot corresponding to the maximum number of extant lineages |
| LTT Slope 2 | slope_2 | Slope of the line connecting the point in the LTT plot corresponding to the maximum number of extant lineages and its endpoint. |
| LTT Slope ratio | slope_ratio | Ratio of LTT Slope 1 and LTT Slope 2. |
| Mean sampling interval | mean_s_time | Mean time between consecutive sampling events, i.e. downward steps in the LTT plot. |
| Mean branching interval | mean_b_time.x | Mean time between consecutive branching events, i.e. upward steps in the LTT plot. This is calculated over three different time periods*. |
| Colless index | colless | Measure of tree imbalance. It is defined as the difference between the numbers of leaves in the left and right subtrees starting from an internal node, summed |

| | | |
|---|---|---|
| | | over all internal nodes. |
| Sackin index | sackin | Measure of tree imbalance. It is defined as the sum of the depths** of all leaves. |
| Maximum ladder size | max_ladder | Size of the largest ladder motif, divided by the number of leaves. A ladder motif is a chain of internal nodes each having exactly one leaf child. |
| Proportion of ladders | IL_nodes | Sum of sizes of ladders, divided by the number of internal nodes. |
| Staircaseness 1 | staircaseness_1 | Proportion of internal nodes with more leaves on one subtree. |
| Staircaseness 2 | staircaseness_2 | Mean ratio of the minimum and maximum numbers of leaves on the left and right subtrees across all internal nodes. |
| Width ratio | WD_ratio | Ratio between the maximum width and the maximum depth*** of a tree. The width count $W_k$ is the number of nodes in the tree whose depth is $k$. The width of a tree is the distance $k$ corresponding to the largest width count. |
| Width absolute difference | $\Delta W$ | It is calculated as the sum of absolute differences between consecutive width counts, i.e. $\sum_k \|W_{k+1} - W_k\|$. |
| LTT coordinates | - | It is an interpolated version of the LTT plot. First, 20 equidistant points $u_l$, $l = 1, ..., 20$ are laid out over the range $[0, T]$, where $T$ is the maximum temporal depth of the tree. Second, the number $n_l$ of lineages at time $t = u_l$ is interpolated directly from the LTT plot using the *interp* function from the *numpy* module in *Python*. This statistic thus comprises 40 values. |

* The time interval spanned by the tree is split into 3 intervals of the same length. An internal branch that overlaps with some interval, even if only partially, will be associated with that interval. Please note that a given branch may be associated with multiple intervals. A node belongs to some interval if it falls inside it.

** The Lineage-through-time (LTT) plot is a piecewise constant function denoting the number of extant lineages at any point in time along the tree. The LTT plot increases by 1 after every split in the tree and decreases by 1 after each leaf (a sampling event).
*** The depth of a node is the number of steps in the path connecting the node to the root of the tree.

**Table S4**: **Prior distributions for the BDEI model**. The priors are the same as in [5,6]. Please note that we only sample ρ from its prior to construct training data but we do not infer this parameter.

| **Parameter** | **Description** | **Prior** |
|---|---|---|
| $R_0$ | Basic reproduction number. | Lognormal(0,1.25) |
| $T_0$ | Delay between the occurrence of the primary case and the start of reporting. | Uniform(0 d,92 d) |
| σ | Rate of onset of infectiousness. | Gamma(shape=0.5, rate=0.167 $d^{-1}$), Restricted within the range [0.0385 $d^{-1}$,1 $d^{-1}$] |
| μ | Recovery rate. | Gamma(shape=0.5, rate=0.167 $d^{-1}$), Restricted within the range [0.0385 $d^{-1}$,1 $d^{-1}$] |
| ρ | Sampling ratio. | Beta(70,30) |

#### Supplementary figures

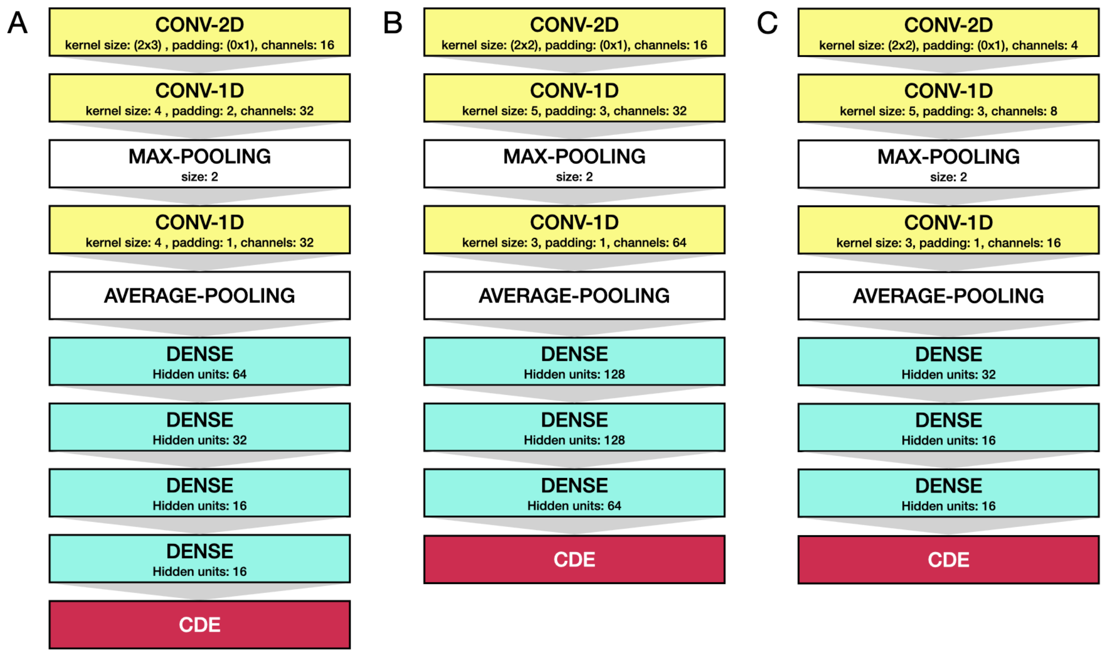


**Fig. S1: Neural network architectures to automatically extract features from raw data**. Panels A to C show the architectures used in the SEIR, BDEI and BD examples, respectively. Each schematics shows to the neural network that maps simulation output $x$ to a low-dimensional vector $y$ according to $y = G_{\phi}(x)$, where $\phi$ denotes the ensemble of neural network parameters. The output $y$ is then fed to the conditional density estimation (CDE) component of the NPE algorithm. Dense blocks consist of artificial neurons that connect to all input features at once. Convolutional blocks' units instead use a small sliding window to perform a 1D or 2D convolution operation on input features. Max- and Average-Pooling layers also use sliding windows to summarise and downscale input signals. Dense and convolutional layers use a RELU activation function [7].

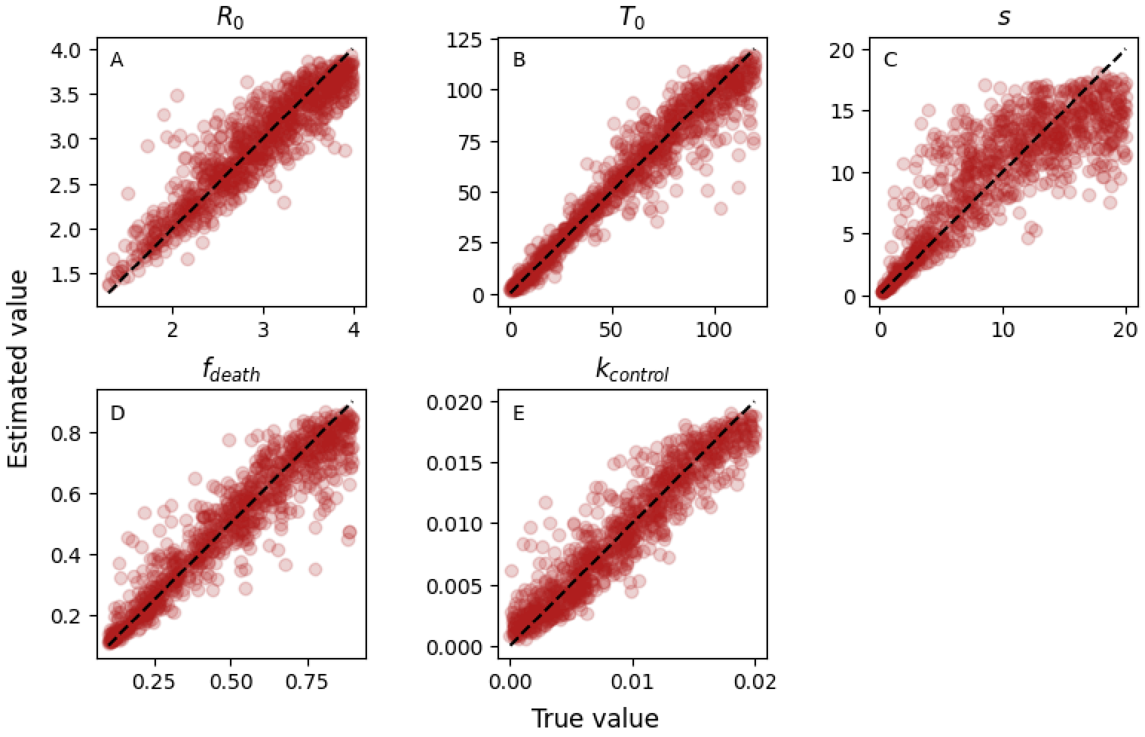


**Fig. S2: Simulation study for the SEIR model.** Estimated vs true parameter values. These results are based on 1000 draws from the joint distribution $p(x, \theta)$.

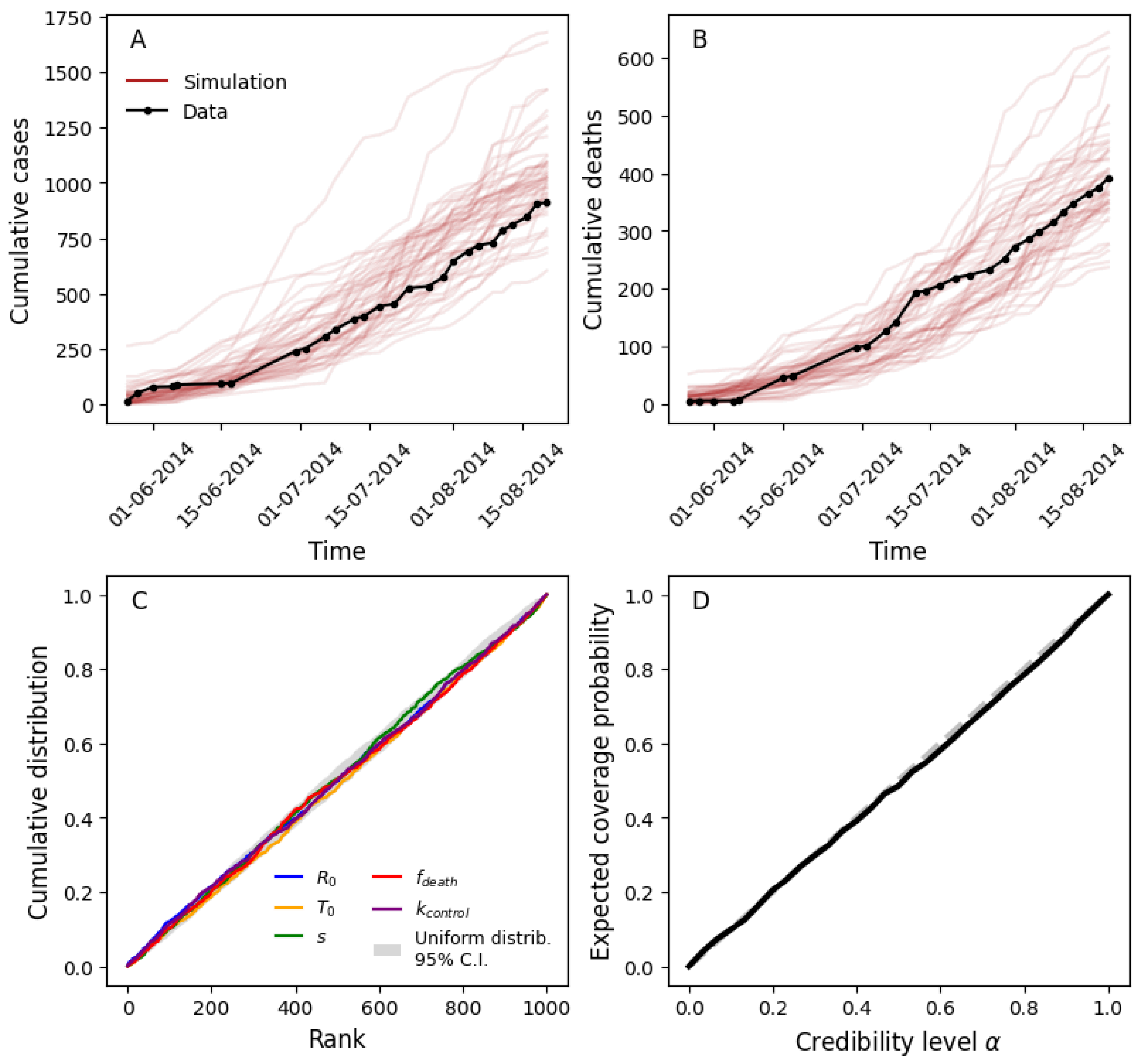


**Fig. S3: Model checking for SEIR model.** (A,B) Red lines represent simulated time series from the posterior predictive distribution $p(x|x_{obs})$ which is conditioned on EBOV data $x_{obs}$ (black). Panels A and B correspond to reported cases and deaths, respectively. (C) Cumulative distribution of ranks for individual parameters as part of rank-based SBC analysis. The latter is based on 1000 draws from the joint $p(x, \theta)$. For each simulation, the rank is calculated using 1000 samples from the estimated posterior. All cumulative distributions fall within the 95% C.I. of the corresponding uniform null distribution. (D) Expected coverage probability vs credibility level as calculated from the TARP analysis. We use 1000 simulations from the joint $p(x, \theta)$ and draw 1000 samples from the estimated posterior for each simulation.

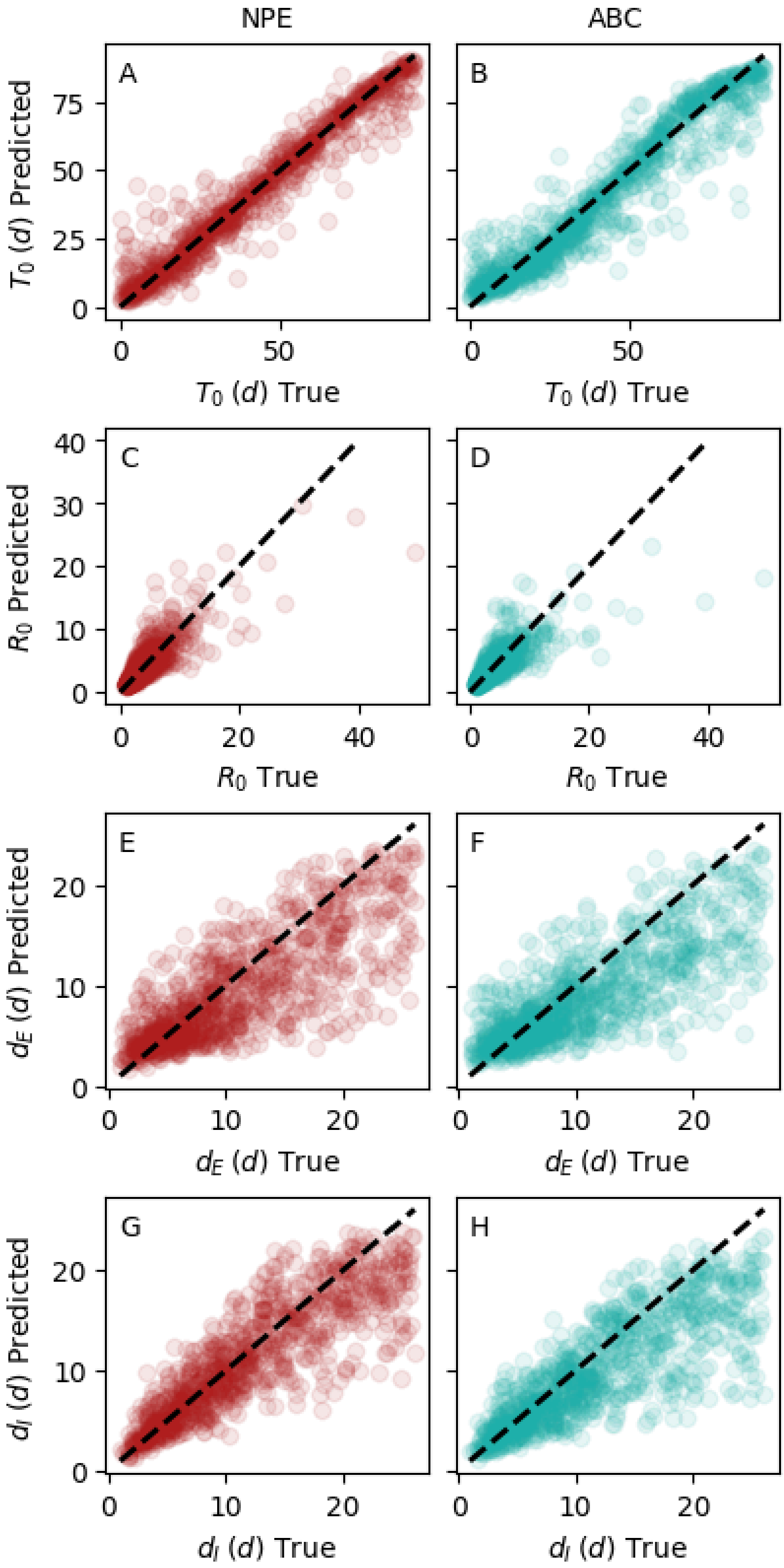


**Fig. S4: Simulation study for the BDEI model.** Estimated vs true parameter values. The left and right columns correspond to NPE and ABC-LR estimates, respectively. These results are based on 1000 draws from the joint distribution $p(x, \theta)$.

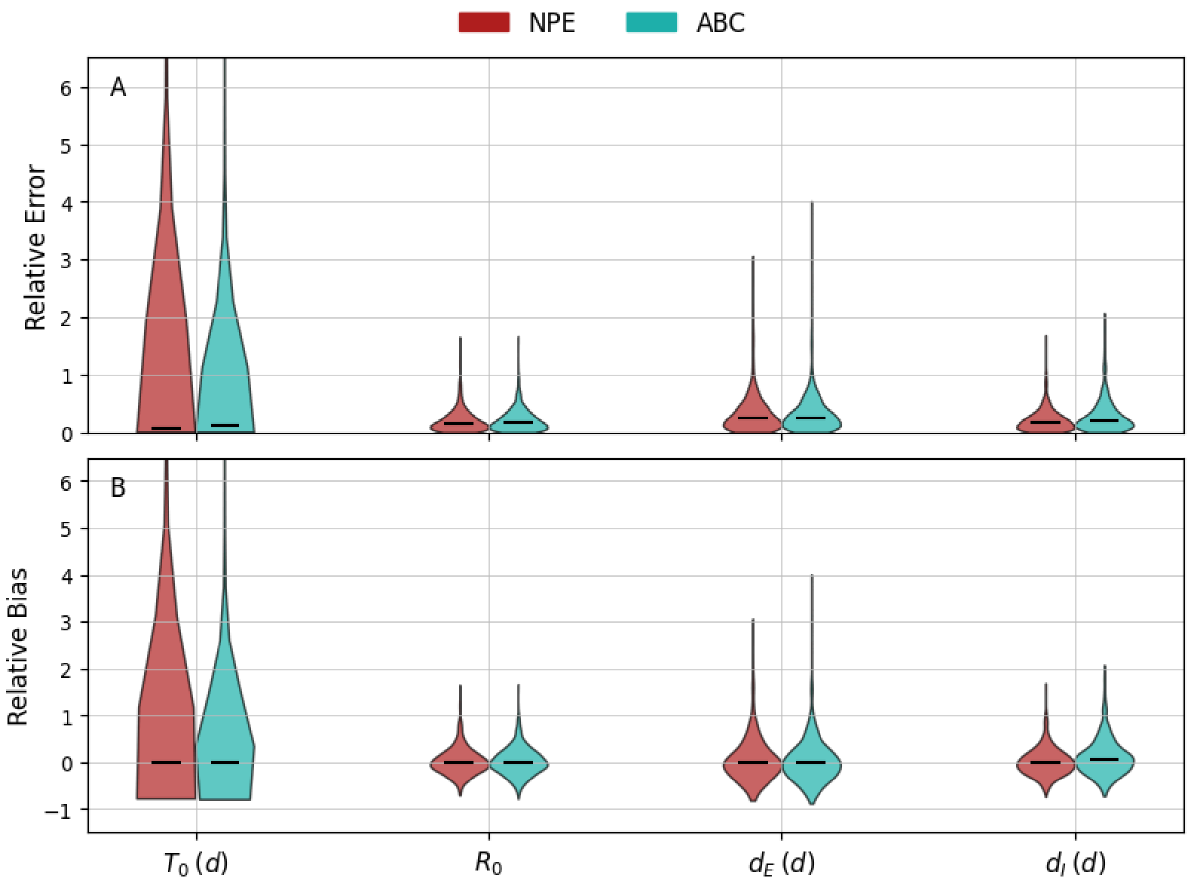


**Fig. S5: Additional calibration metrics for the BDEI model.** (A) Distribution of Relative Error for NPE (red) and ABC-LR (turquoise) for individual parameters. The Relative Error is calculated as $|\hat{\theta} - \theta_{true}|/\theta_{true}$, where $\hat{\theta}$ is the median posterior estimate and $\theta_{true}$ is the true parameter. (B) Distribution of Relative Bias. The Relative Bias is calculated as $(\hat{\theta} - \theta_{true})/\theta_{true}$. Results are based on 1000 simulations from $p(x, \theta)$ and 10000 posterior samples for each simulation. Black bars indicate median values.

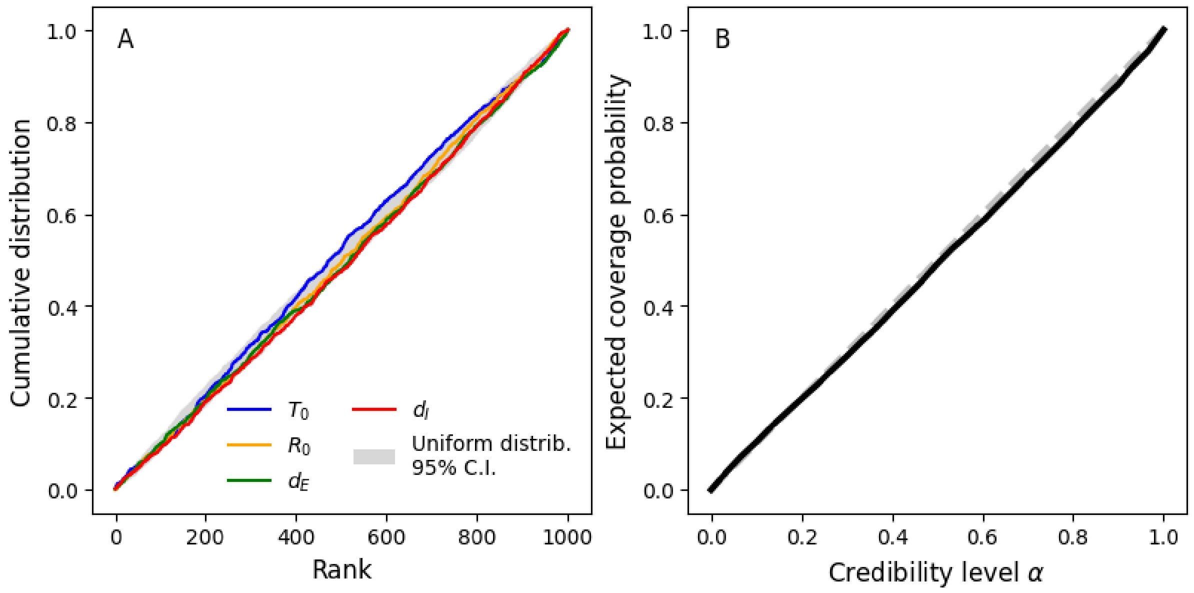

**Fig. S6: Model checking for BDEI model.** (A) Cumulative distribution of ranks for individual parameters as part of rank-based SBC analysis. The latter is based on 1000 draws from the joint $p(x, \theta)$. For each simulation, the rank is calculated using 1000 samples from the estimated posterior. (B) Expected coverage probability vs credibility level as calculated from the TARP analysis. We use 5000 simulations from the joint $p(x, \theta)$ and draw 5000 samples from the estimated posterior for each simulation.

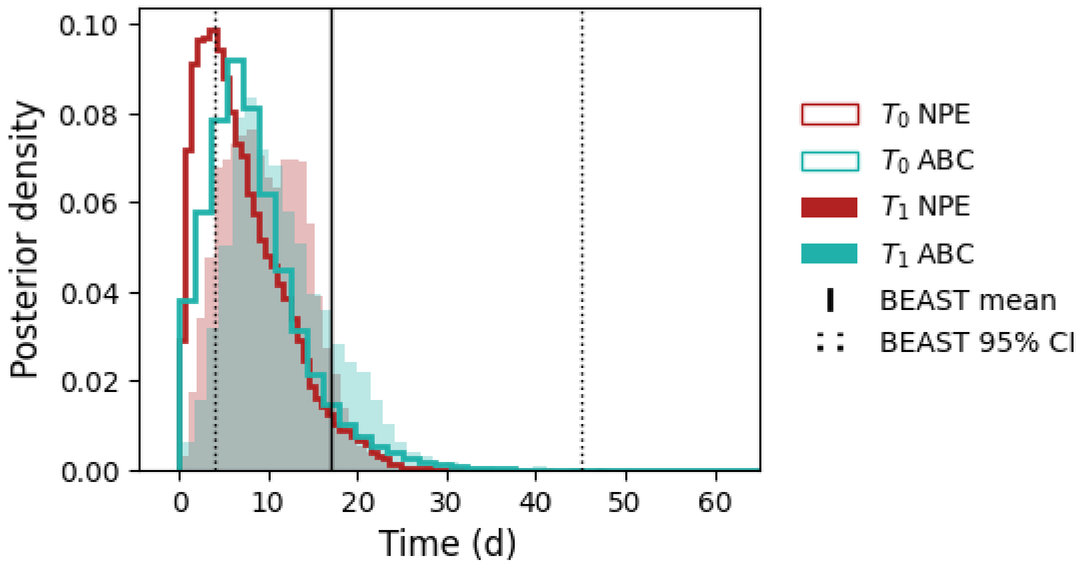


**Fig. S7: Time to first sample.** Here we show the distribution of the delay $T_1$ between the occurrence of the primary case and the first reported sample in the phylodynamics example (filled). $T_1$ is different from $T_0$, which denotes the delay before reporting begins (solid lines, same as in Fig. 2 in the main text). By construction, $T_1 > T_0$. $T_1$ enables a comparison with the reference analysis (black lines), which assumed the sampling ratio to be 0 before the first sample [6]. Our analysis yields a shorter estimate of $T_1$ compared with the original analysis but the corresponding posterior distributions and means are compatible with each other. Results are based on 10000 simulations.

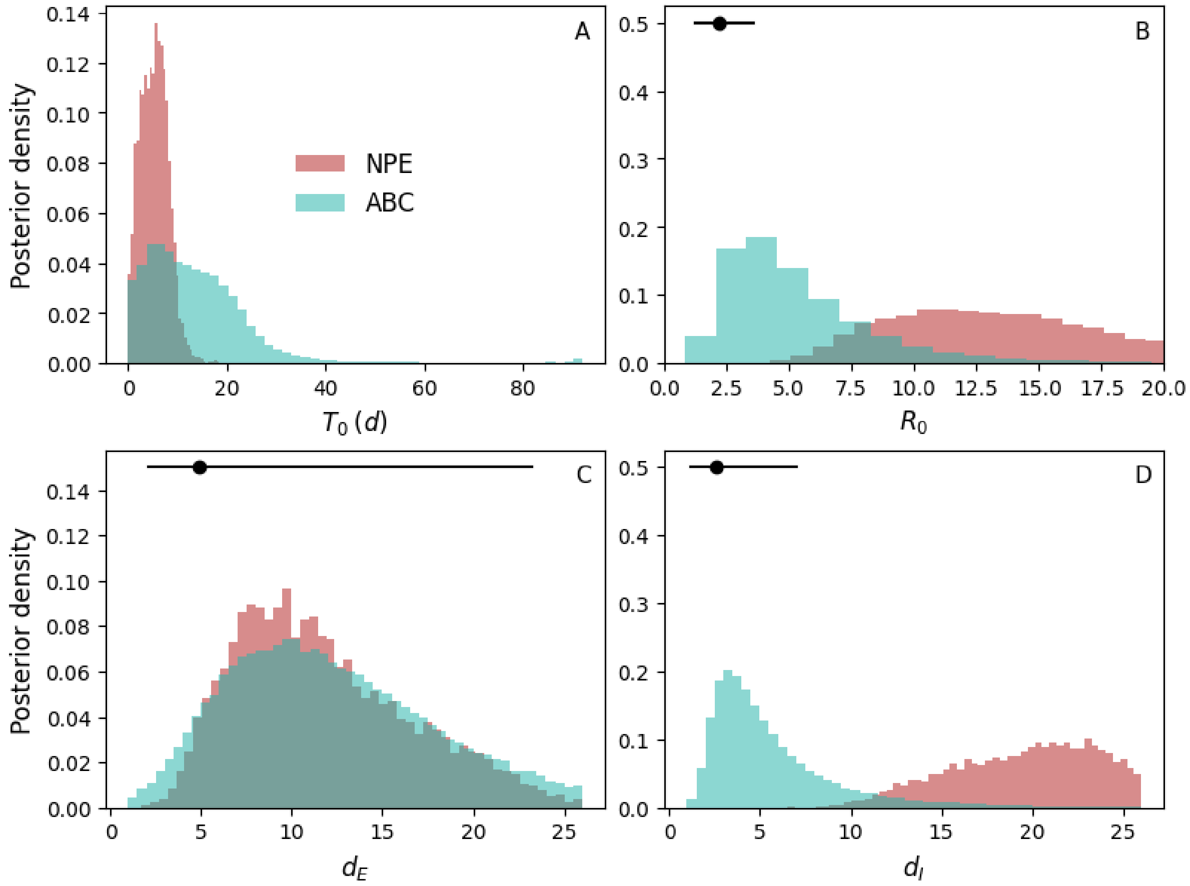


**Fig. S8: Comparison of NPE and ABC-LR inference results on EBOV MCC tree.** NPE and ABC-LR yield different predictions on the MCC tree. Importantly, both methods yield parameter estimates that are not compatible with original MCMC estimates (black dot: mean, black line: 95% C.I.). The MCC tree is not an appropriate summary of the posterior distribution over trees (see Fig. S9).

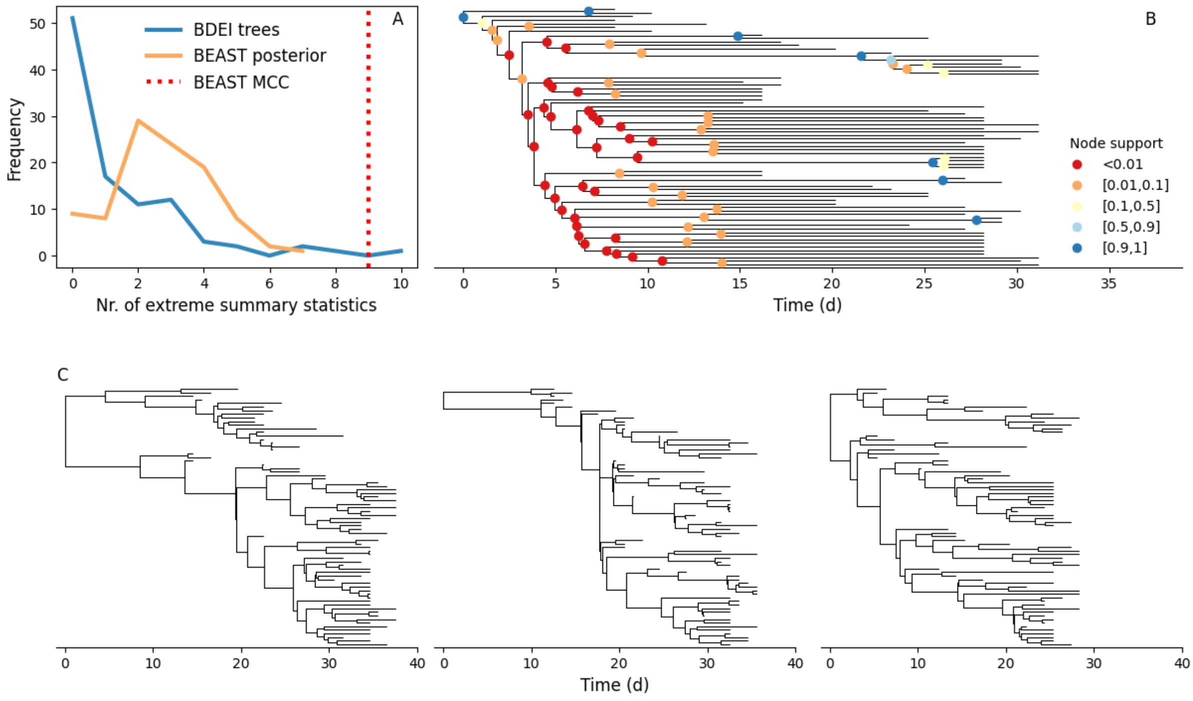


**Fig. S9: The MCC tree is not representative of posterior or simulated trees.** (A) Distributions of counts of extreme summary statistics per tree for different tree sets. We calculated the distributions of 31 summary statistics (Those retained after VIF selection) from 200000 BDEI trees drawn from the joint distribution $p(x, \theta)$. A summary derived from some tree is said to be extreme if it falls outside the 95% C.I. of this distribution. The blue line shows counts from 100 trees drawn from the joint $p(x, \theta)$ and acts as a reference expectation. The orange line corresponds to the 100 BEAST posterior trees used in Fig. 2 in the main text. The red line denotes the number of extreme summary statistics for the MCC tree. This figure shows that the MCC tree is an outlier that is poorly represented by BDEI simulations. Posterior trees display some extreme features as well, though not nearly as many as for the MCC tree.The MCC and a few posterior trees are shown in (B) and (C), respectively. Nodes in the MCC tree are painted according to their posterior support. Most internal nodes are poorly supported.

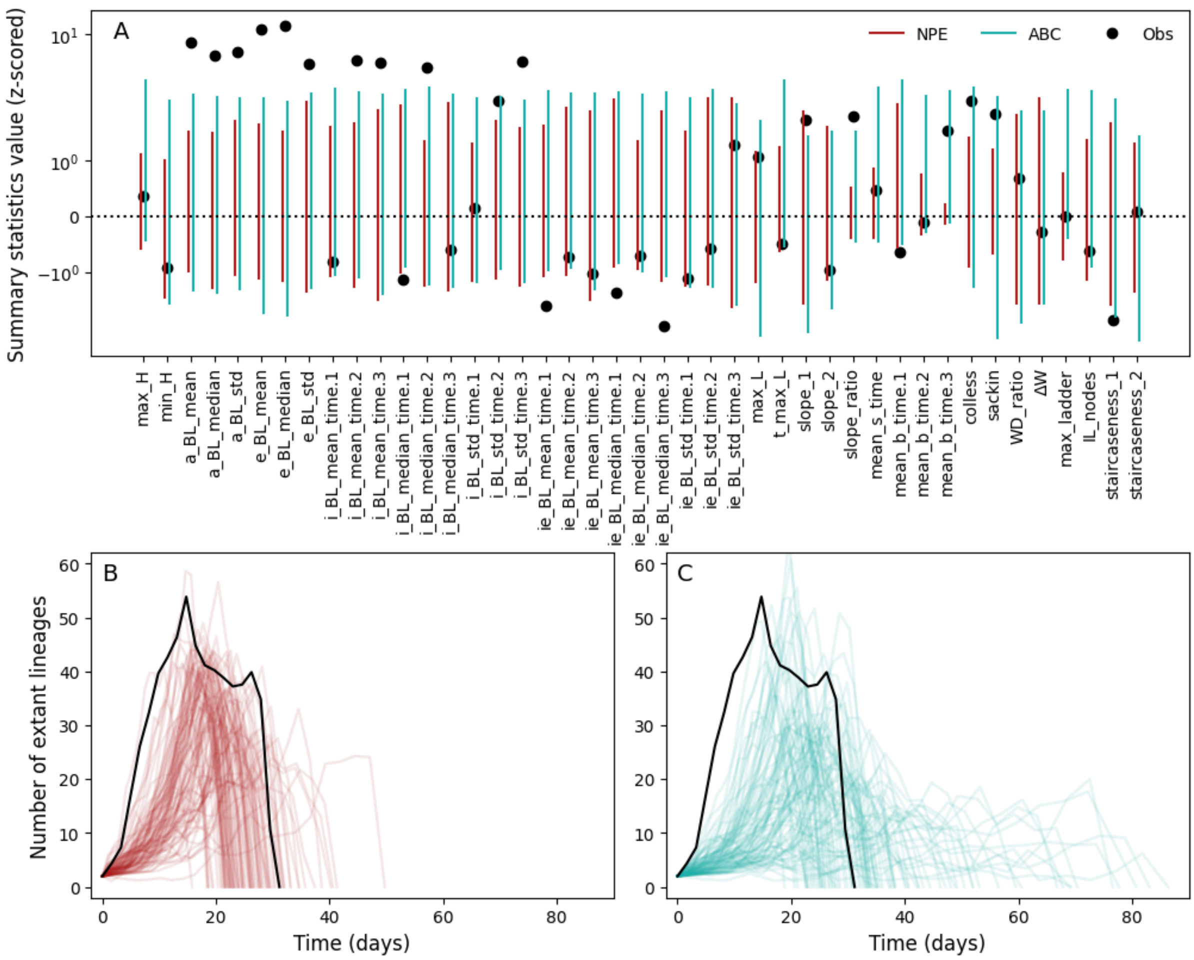


**Fig. S10: NPE and ABC-LR fail posterior predictive checks when considering the MCC tree.** (A) Each bar represents the 95% C.I. for a given summary statistic (described in Table S3) for either the NPE (red) or the ABC-LR (teal) fits. Black dots represent the summary statistics calculated from the MCC tree. All values were z-scored based on the mean and standard deviation calculated from the NPE posterior predictive simulations. Results are based on 200 simulations from the posterior predictive distribution. (B,C) Comparison of LTT plots calculated from the MCC (black) and posterior predictive trees (see LTT coordinates summary statistics in Table S3). The NPE fit is shown in (B) while the ABC-LR fit is shown in (C). Coloured lines represent 100 independent posterior predictive simulations.
